# Preparing future STEM faculty nationwide through flexible teaching professional development

**DOI:** 10.1101/2022.10.06.511090

**Authors:** B. B. Goldberg, D. Bruff, R. Greenler, K. Barnicle, N. Green, L. E. P. Campbell, S. L. Laursen, M. Ford, A. Serafini, C. Mack, T. Carley, C. Maimone, H. Campa

## Abstract

We report on a five-year initiative that has prepared thousands of future STEM faculty around the world to adopt evidence-based instructional practices by participating in two massive open online courses (MOOCs) and facilitated in-person learning communities. This novel combination of asynchronous online and coordinated, structured face-to-face learning community experiences provides flexible options for STEM graduate students and postdoctoral fellows to pursue teaching professional development, while leveraging the affordances of educational technologies and the geographically clustered nature of this target learner demographic. A total of 14,977 participants enrolled in seven offerings of the introductory course held 2014-2018, with 1,725 participants from approximately 60 countries completing at an average course completion rate of 13%. The preparation of future STEM faculty makes an important difference in establishing high-quality instruction that meets the diverse needs of all undergraduate students, and the initiative described here can serve as a model for increasing access to such preparation.

## Introduction

There is recognition [1] that evidence-based, student-centered instruction in science, technology, engineering, and mathematics (STEM) increases undergraduate student learning and success in STEM generally [2] and reduces the performance disparities between majority and minority students in STEM [3, 4, 5, 6]. There is also evidence that current [7] and future [8] faculty who engage in effective professional development go on to implement evidence-based pedagogies in their classes. These findings are the basis for pedagogical professional development programs offered by university teaching centers, graduate schools, and postdoctoral training initiatives [9].

Future STEM faculty, that is, doctoral students and postdocs who seek academic careers, face particular challenges in learning about and adopting evidence-based teaching practices, including limited opportunities and lack of advisor support for pedagogical professional development [10, 11, 12, 13]. Despite this, graduate students and postdocs may be more receptive than current faculty to explore and implement evidence-based teaching practices because they are in the process of learning the standards of academia, developing scientific and teaching practices in their discipline, and are preparing for competitive academic positions [14]. Encouragingly, future STEM faculty who do participate in moderate- or high-engagement pedagogical professional development (greater than 25 hours of participation) report significantly improved self-efficacy as instructors and significantly higher adoption of evidence-based teaching practices [8] and perform as well or better in research [15].

To provide such professional development to future STEM faculty, and thereby improve undergraduate education in the U.S. more broadly, the Center for the Integration of Research, Teaching, and Learning (CIRTL) Network, which currently consists of 43 research universities across the United States and Canada, provides structured pedagogical professional development programs for graduate students and postdocs at individual campuses and through cross-Network programming [16, 17, 18]. Many of these programs are structured as in-person or virtual, synchronous or asynchronous learning communities [19, 17], where participants meet to learn from and with each other as they pursue shared learning goals [20]. The Network also serves as a community of practice [21] for leaders of future STEM faculty development to share strategies and expertise and to co-develop and implement network-wide programs.

In 2013, a series of CIRTL Network conversations on emerging models for future STEM faculty development led a small group of faculty, administrators, and researchers to propose a new initiative centered on the use of Massive, Open Online Courses (MOOC). Interest in this new form of online education accelerated rapidly [22], with educators and researchers exploring the potential for online tools such as videos, discussions, and peer assessments to support learning for thousands of concurrent students [23]. Interestingly, research shows that the more successful MOOCs have been associated with targeted rather than general audiences [24].

In this context of pedagogical experimentation and with funding from the National Science Foundation, we sought to design, deliver, and evaluate the use of MOOCs on evidence-based undergraduate STEM teaching for future faculty pedagogical professional development. This in itself was not novel; other MOOCs developed in the same time frame also had this focus [25].

Inspired by instructors who “wrapped” campus-based courses around existing MOOCs [26] and informed by the CIRTL Network’s experience with campus-based and virtual learning communities, we planned the online courses to be delivered in three different modes to meet the diverse learning needs of future faculty : (1) as stand-alone MOOCs for online participants, (2) as open educational resources for use by individuals or by campus-based professional development programs, and (3) as blended online and in-person experiences constructed with what we called MOOC-Centered Learning Communities, or MCLCs [27]. By inviting colleagues around the CIRTL Network and beyond to host MCLCs of participants in the online courses and by providing those local facilitators with learning guides to support their local in-person meetings, we designed a novel structure that has enabled us to meet the professional development needs of thousands of future STEM faculty worldwide.

## Materials and methods

In 2013-2014, we launched an eight-week introductory MOOC, *Introduction to Evidence-based Undergraduate STEM Teaching*, followed by a second eight-week MOOC, *Advanced Learning Through Evidence-Based STEM Teaching*, in 2015-2016. Each course consists of six 3-to-5-hour modules, each featuring instructional videos, discussion prompts, recommended readings, and a quiz, and three peer-graded assessments (PGAs) per course. The introductory course examines the fundamentals of learning and learning design, including learning objectives, assessment, and active learning, culminating with a final PGA in which participants develop a sample lesson plan incorporating these core elements. The advanced course delves deeper into evidence-based teaching practices, including peer instruction, cooperative learning, and inquiry-based labs. The final PGA in the second course requires participants to develop a teaching philosophy statement that demonstrates their understanding of and preferences among the teaching practices they have learned. Each course is offered once or twice a year on the edX platform.

To foster greater engagement and learning, we encouraged participants in the online courses to join or start an MCLC. These learning communities are typically hosted on university campuses, and typically meet weekly to share, discuss, and contextualize what participants are learning in the online course. Depending on local needs, MCLCs can be part of credit-bearing courses, non-credit seminars, or an informal set of meetings among peers or colleagues. Each MCLC has a facilitator who regularly convenes the community and plans discussions or other activities for the in-person meetings. To support MCLC facilitators, we provided an “MCLC Facilitators’ Guide” that includes learning goals and objectives for online and in-person sessions; overviews of online videos, discussion prompts to engage participants with course content, and assignments; and 3-7 suggested activities with facilitator notes for each module that complement and extend the online materials. Our project team markets the potential of MCLCs for professional development associated with teaching and learning and recruits MCLC facilitators at CIRTL Network campuses as well as through our respective networks to draw a diverse and international community.

The project website, https://www.stemteachingcourse.org/, makes freely available the course content, including videos, accompanying slides, discussion prompts, and instructions for each PGA. The project also has a public YouTube channel, https://www.youtube.com/user/cirtlmooc, featuring all course videos organized by course module. All materials are made available under a Creative Commons 4.0 Attribution-Noncommercial license to facilitate reuse by anyone interested in STEM teaching or pedagogical professional development.

## Results

### Outcomes of participation and engagement

To understand the impact of the project, we examined the learner experience: Who engaged in the course, how they engaged, what motivated them, and what they thought of it. A total of 14,977 participants enrolled in seven offerings of the introductory course held 2014-2018, with 1,725 participants from approximately 60 countries completing. The average course completion rate of 13% is more than double the rate of most non-professional and non-degree MOOCs [28, 29, 24].

Overall, 5,320 total participants registered for the four offerings to date of the second MOOC, with 291 completers and a course-averaged 6.3% completion rate (Table 1).

**Table 1:**
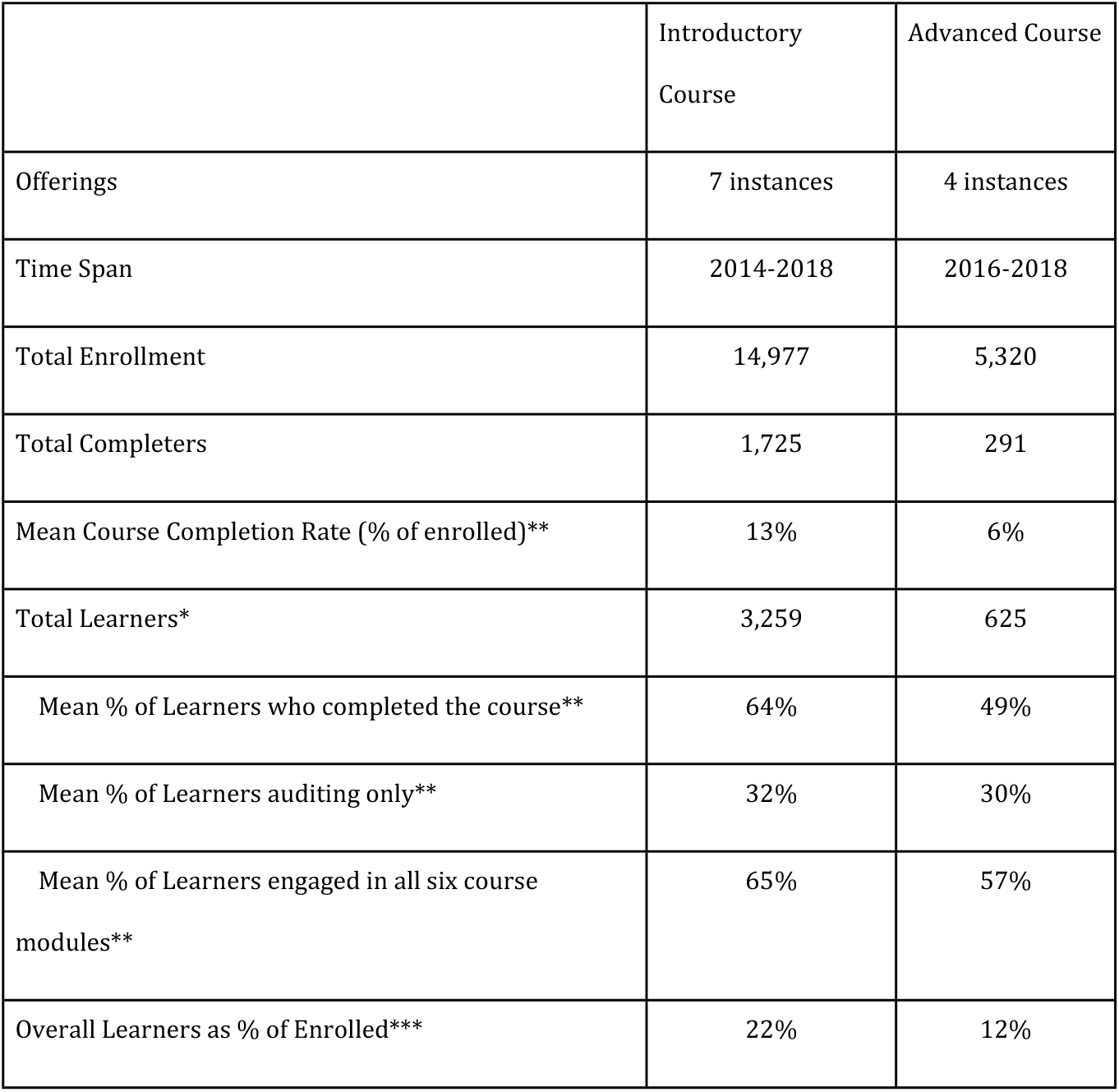
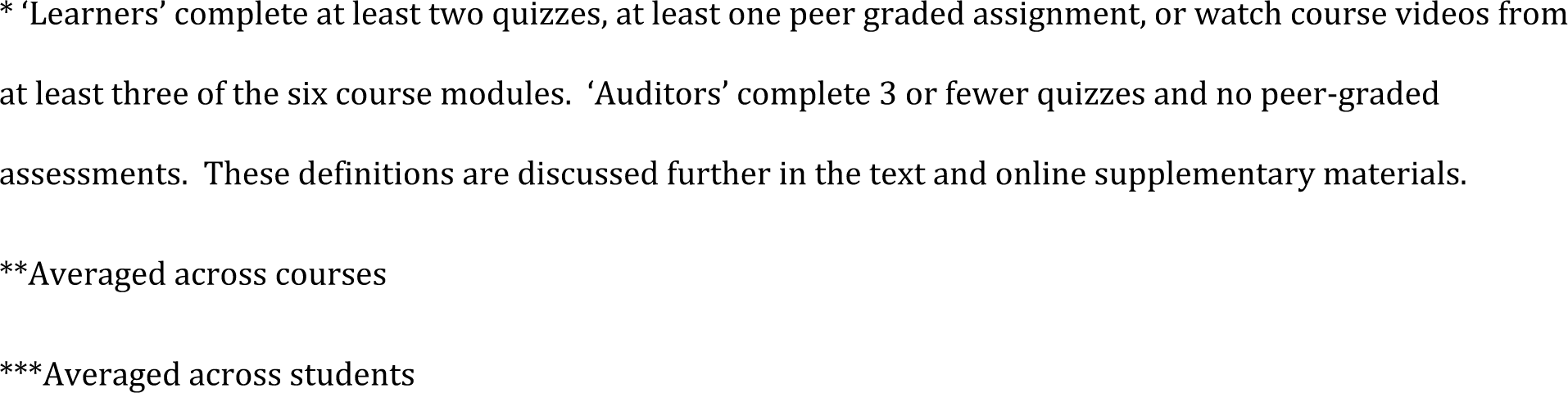
Participation, learner engagement and completion for 11 course offerings from 2014-2018.

|  | Introductory Course | Advanced Course |
| --- | --- | --- |
| Offerings | 7 instances | 4 instances |
| Time Span | 2014-2018 | 2016-2018 |
| Total Enrollment | 14,977 | 5,320 |
| Total Completers | 1,725 | 291 |
| Mean Course Completion Rate (% of enrolled)** | 13% | 6% |
| Total Learners* | 3,259 | 625 |
| Mean % of Learners who completed the course** | 64% | 49% |
| Mean % of Learners auditing only** | 32% | 30% |
| Mean % of Learners engaged in all six course modules** | 65% | 57% |
| Overall Learners as % of Enrolled*** | 22% | 12% |

\* ‘Learners’ complete at least two quizzes, at least one peer graded assignment, or watch course videos from at least three of the six course modules. ‘Auditors’ complete 3 or fewer quizzes and no peer-graded assessments. These definitions are discussed further in the text and online supplementary materials.
\*\*Averaged across courses
\*\*\*Averaged across students

We analyzed participants’ course activity, engagement, and outcomes in the introductory MOOC based on data from the online course platforms, and voluntary pre- and post-course surveys [30]. Further analysis is available in the online supplementary materials.

Course completion required participants to pass quizzes, PGAs, or a combination thereof. These assessments required understanding of and ability to apply the material from the course videos and readings. While completion is one marker of participant engagement in MOOCs, we encouraged participants to make use of the course materials in whatever manner would best support their development. In assessing engagement, we focused on those who engaged in significant ways with the course materials, as a proxy for learning and gains in knowledge. We define “learners” to be participants who complete at least two quizzes, complete a PGA, or watch videos from at least three of the six course modules. These thresholds are based on drop-offs observed in participant activity as a function of both video watching and assignment completion (Fig 1a). (Note that the course included a module “0”, introducing the course.) As others have also found, course engagement drops off quickly after the first and second weeks [31, 32].

**Fig 1:**
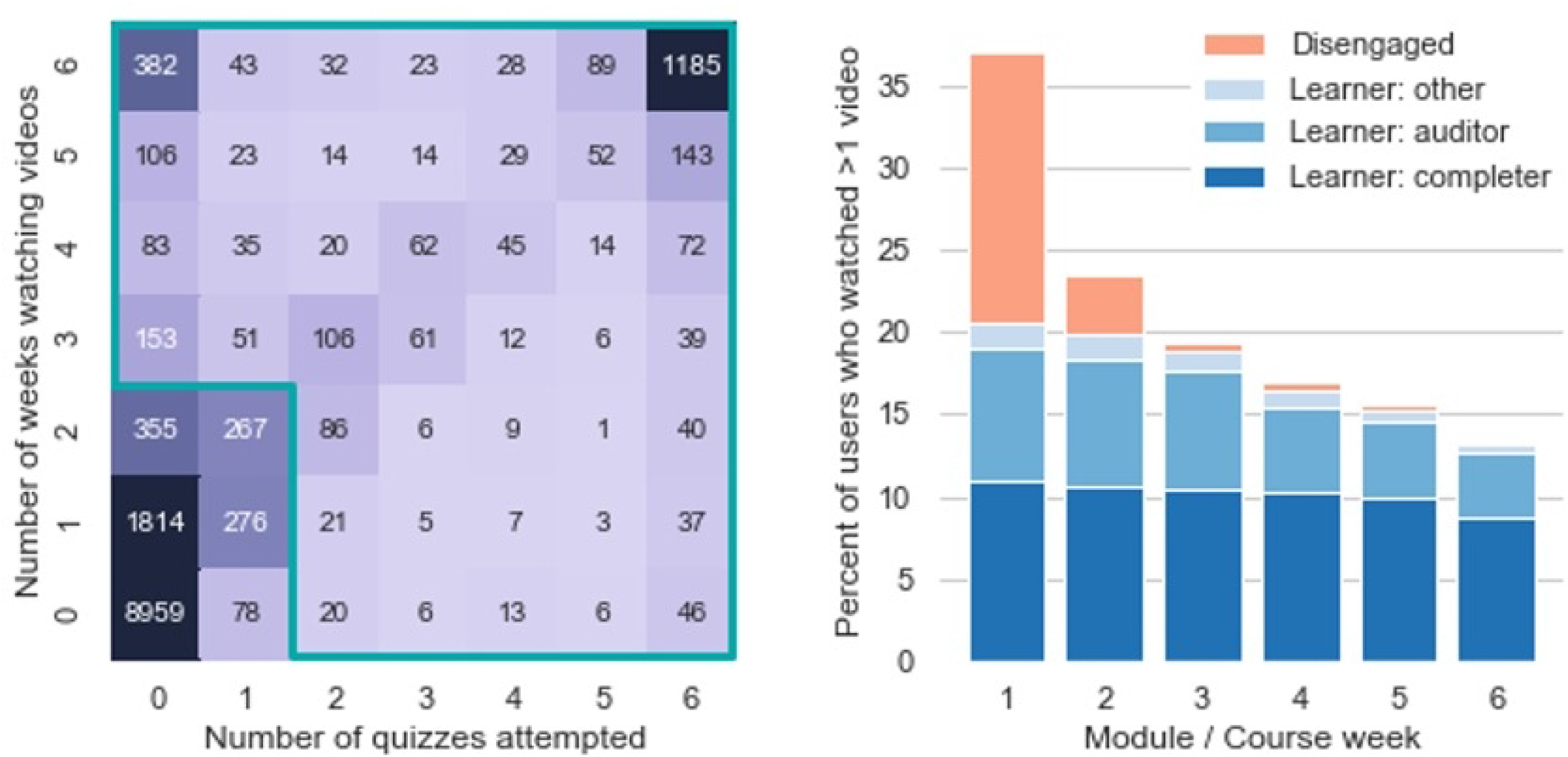
Learner engagement v. video watching, assignments, and weeks. **Fig 1 (a)** Joint histogram of enrolled in the introductory MOOC by total number of modules/course weeks in which they participated by watching videos (vertical axis) or taking quizzes (horizontal axis). The outlined region approximately separates “learners” from disengaged non-learners; a small number (31 or 1%) of learners who completed a PGA but few quizzes may not fall within the outlined region. **Fig 1 (b)** Percent of total enrolled who participated during each module/course week by watching more than one video that week.

Learners represented 22% of those enrolled in the introductory course. Learners fell into two main groups: completers (64% of learners, or 14% of all enrolled) who participate in quizzes and peer-graded assignments, and auditors (our term, representing 32% of learners, 7% of enrolled) who primarily watch course videos (see [33]). As seen in Fig 1(b), non-learners tended to be active in the first two weeks of the course, but later disengaged from the course. About half the auditors did not meet the course completion criteria but still participated in all six of the course modules by watching a module video or taking the module quiz. Thus, 78% of the learners engaged continuously with the material throughout the course, indicating a high rate of retention beyond the first two weeks.

Voluntary pre- and post-course surveys inquired about intention to complete the course, learning gains, motivations, time spent, involvement in learning communities, demographics and other information about our participants, their behaviors and learning outcomes. Demographic data about course participants is only available through these surveys.

For seven instances of the introductory course, 3,884 students (26% of enrolled) took the pre-course survey. In a subset of the data where we can link participant demographics to course engagement behaviors, pre-course survey respondents included 57% of learners in the course; conversely, 55% of pre-course survey respondents went on to engage with the course as learners. This overlap implies that pre-course survey respondents are a very good, but not perfect, representation of learners. Similarly, at the end of the course, half (55%) of the completers responded to the post-course survey; among post-survey respondents, 84% completed the course and an additional 8% engaged during all six modules/weeks. Results from the post-course survey are, therefore, very reflective of the experiences and demographics of course completers.

PhD students and postdocs made up 50% of the pre-course survey respondents and 59% of the post-course survey respondents who indicated their status (98.8% and 90.3% respectively), representing by far the largest audience segment of both learners and completers. Faculty made up an additional 20% and 19% of the pre- and post-course survey respondents respectively. Nearly all (91%) of the pre-course survey respondents indicated their disciplines from STEM or Social, Behavioral, and Economic Sciences (SBES) fields. Thus, we are reaching our designed audience of STEM PhDs and postdocs, 86% of whom reported preparing for academic careers.

Overall, 34% of pre-course survey respondents completed the course. Among them, postdoctoral researchers completed at a higher rate of 39% compared to 32% for other participants (p = 0.005). Those who indicated on the pre-course survey that they intended to pursue an academic career (74% of respondents) completed at a significantly higher rate, 37%, than those who did not, 23% (p << 0.001).

Post-course survey respondents rated their retrospective gains as high in four areas: *confidence of implementing teaching and learning strategies covered in class, interest in taking or planning to take additional classes related to teaching and learning, interest in discussing teaching and learning with colleagues and friends*, and *confidence that they understand the material covered* all as higher than 4.0 on a 5-point scale where 5 is “*Great gains*.*”* Participants who responded to both pre- and post-course surveys also reported increased familiarity with the key concepts of summative assessment, leveraging diversity, formative assessment, backwards design, and learning objectives that were taught in the course (Fig 2).

**Fig 2:**
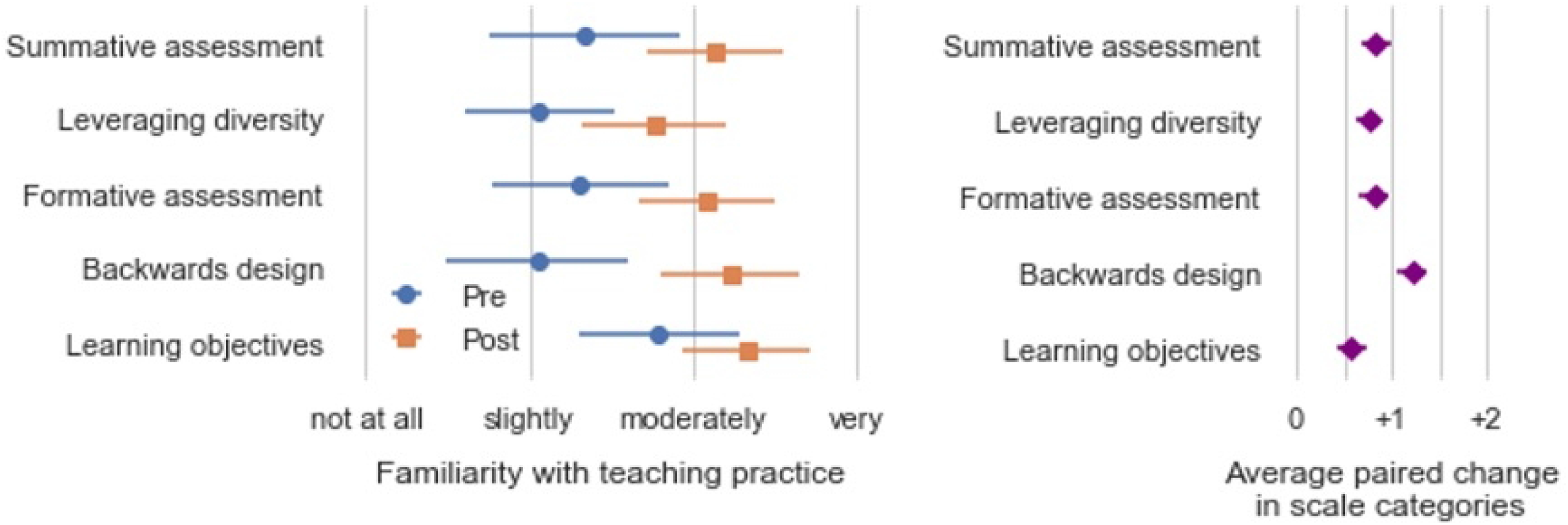
Increase in average reported familiarity with pedagogical topics from course participants. **Fig 2 (a)** Average responses of pre- and post-course respondents, unpaired. Error bars represent one standard deviation. **Fig 2 (b)** Average of paired differences for the 520 respondents who took both the pre- and post-course surveys for the first two instances of the course where responses can be linked. Error bars represent the 99% confidence interval.

The high rate of learner completion, especially among those who self-identified as future STEM faculty, indicated strong motivation of participants and a course design that matches learners’ time commitment, work level, availability and motivation. These conclusions are corroborated by self-reported satisfaction of post-course survey respondents, 97% of whom agreed that the course improved their ability to teach, 93% were either “satisfied” or “extremely satisfied” with the course, and 97% would “recommend [the course] to others” [34].

### Learning community engagement

Many learners engaged in our blended model of delivery: 134 institutions have hosted at least one in-person learning community, and many have hosted multiple times, yielding 236 total MCLCs as of Spring 2018. MCLCs had on average 12 participants, who were largely (75%) STEM PhD graduate students and postdocs, and who completed the course at a high rate (65%), as estimated by MCLC facilitators.

We do not have data on how many learners were in MCLCs. Based on data about their intentions [35], our best estimate is that between 20-40% of learners were in MCLCs. The top reasons they wished to engage in local, in-person learning were the opportunities to *interact with peers* (35%), to *discuss course materials and assignments* (32%), to *meet others interested in teaching and learning* (31%), and to *receive feedback on my teaching and learning practices* (28%). Among post-course survey respondents, 34% reported participating in an MCLC, and those who did had strong outcomes: 97% were learners and 87% completed the course, representing 19% of all completers.

### Feedback from learning community facilitators

Survey responses and interviews with MCLC facilitators provided valuable feedback on the structure and efficacy of the MOOCs broadly and the MCLCs in particular. Their thoughtful feedback informed substantial revisions of the introductory and advanced MOOCs. In their MCLCs, facilitators reported using the facilitator guides and finding most components to be useful (∼75%); they reviewed it to get ideas and used different activities to meet the needs of their particular group. Activities involving reflection, discussion or extension of course material were well received, while those that relied on participants’ past teaching experience, or required peer feedback, additional reading, or reflection outside the MCLC meeting, were generally less successful. Both facilitators with prior expertise in the MOOC content and those without prior knowledge reported success in leading MCLCs: experts tended to prepare MCLCs as mini-courses enriched with their own content and activities, while novices conducted MCLCs in the form of peer-led study groups, largely drawing on the MOOC materials and the facilitator guide. That novice and expert leaders can successfully lead MCLCs with the support of the Guide, makes the MCLC model sustainable and adaptable in numerous settings.

Most (45 of 51) facilitator survey respondents reported they would facilitate an MCLC again, saying, for example, *“I enjoyed facilitating the MOOC, learning from it, and sharing my experience with the participants in our learning community,”* and “*It’s one of my favorite things to do, even though I am doing it as a volunteer*.”

### Open educational resource engagement

In addition to the stand-alone MOOC and MCLC delivery modes, open educational resources (OER) are intended to encourage adoption and adaptation broadly and done in the spirit of collaboration inherent in evidence-based teaching strategies. By making access as broad and simple as possible, we made the tradeoff of limiting our ability to monitor participants and facilitators who have accessed our content. Multiple colleagues and educators expressed an interest in using our course materials for professional development programs at their institutions. In the first three years our ∼130 videos were viewed 60,540 times outside the course. Examples of how educators have used our materials include: developing or supporting teaching certificate programs, redesigning curricula, incorporating additional materials into existing educational workshops or for credit courses, and providing online professional development training.

## Discussion

Through multiple offerings of two MOOCs on evidence-based undergraduate STEM teaching, intentional support for facilitated MCLCs, and open access to course materials, we have met a need among graduate students and postdocs for pedagogical professional development that often goes unmet through traditional on-campus resources and events [36]. A number of key factors led to this result.

Our target audience of STEM graduate students and postdocs have clearly identified professional development goals and are geographically clustered at research universities. This enabled the formation of local MCLCs, since potential participants studied and worked in proximity to each other. Having local MCLCs at universities also made easier publicity and recruitment for our blended delivery mode, since the opportunity to join MCLCs could be advertised by supportive university faculty and staff members, graduate schools, departments and centers for learning and teaching. Those faculty and staff also made ready facilitators for MCLCs. Their self-reported experience and expertise, and the similar professional goals of participants, lent MCLCs a structure and coherence that distinguished them from the ad hoc student meet-ups that are common in many MOOCs, leading to greater course completion rates by MCLC participants.

Participants had flexible options for engaging with course materials and resources. Some learners completed the courses by submitting quizzes and peer-graded assignments, some audited the courses by consistently watching videos, while other learners viewed course videos in an ad hoc manner on YouTube and the project website. MCLCs provided a professional development option for those who also wanted an in-person experience. For graduate students and postdocs often constrained by time, advisor priorities and the need to focus on research, these options enabled motivated future faculty to seek out and obtain pedagogical professional development on their own terms.

Our MOOC initiative was launched from, developed by, and continues to be hosted by an existing network of STEM faculty, educators, administrators, and educational developers, the CIRTL Network. Long-term sustainment is an unfortunately rare outcome of NSF-funded educational initiatives [37]. The CIRTL Network brought together the initial team that developed the MOOCs; it provided a range of STEM education practitioners and researchers who contributed to the course content through interviews, resource sharing, module development, and feedback; and Network institutions hosted approximately one third of the MCLCs. Individuals from outside the CIRTL Network contributed in significant ways as well, particularly as MCLC facilitators, but the existing network functioned as a community of practice that enabled the initiative to succeed. From 2018 to now, the CIRTL Network has assumed all management of the MOOCs and MCLCs, with continued success.

Recent research shows that, while MOOC participation and completion rates have declined over the last five years, MOOCs designed for highly motivated students pursuing professional development have thrived [24]. Our findings are consistent with this trend and point to potential future uses of MOOCs and MCLCs for career and professional development needs. Asynchronous, online learning in conjunction with synchronous, in-person learning is a structure with potential to be effective in professional development domains beyond teaching, including leadership, conflict resolution, responsible conduct of research, and mentoring, as well as interdisciplinary domains such as data visualization or computational thinking. The fact that current STEM faculty also took our MOOCs and participated in MCLCs suggests that this structure might also be useful for early-career academics especially at institutions without faculty development programs, as long as they are structurally and geographically clustered. In this project, the CIRTL Network was instrumental, but other professional networks such as disciplinary societies, formal and informal, could serve similar design, support, dissemination, and sustainment functions.

We demonstrated the effective delivery of pedagogical professional development to future STEM faculty with the potential to significantly impact undergraduate STEM education across the United States. Our design combines flexible, asynchronous content in conjunction with optional, supported and facilitated in-person learning communities, all within the context of a national network of STEM faculty and educational developers. Our model can successfully be used in many contexts to overcome barriers where learners seek significant professional development in constrained settings.

## Acknowledgments

We gratefully acknowledge our many colleagues that contributed to this effort by developing MOOC modules or content to components of modules. We specifically would like to thank (in alphabetical order): C. Brame, S. Chasteen, M. DiPietro, C. Fata-Hartley, A. Little, J. Littrell, R. M. Mathieu, T. McMahon, and K. Spilios, for their many critical contributions.

## Author contributions

Proposed and secured financial support (BBG, DB, HC, KB); developed MOOC content and structure (LEPC, CM, BBG, DB, HC, KB, NHG, RG, TC), built and/or managed courses on edX and Coursera (LEPC, AS, BBG, DB, HC, KB, NHG, RG, TC), prepared MCLC Facilitators guides (BBG, DB, HC, KB, NHG, RG), analyses (SL, MF, CM, AS, BBG, DB, HC, KB, NHG, RG), prepared the manuscript (SL, MF, CM, AS, BBG, DB, HC, KB, NHG, RG)

SLL interviewed MCLC and institutional leaders, analyzed data, and contributed to the manuscript.

### One sentence

We’ve prepared tens of thousands of future STEM faculty to use evidence-based instruction through a MOOC in conjunction with hundreds of supported and facilitated in-person (now virtual) learning communities that provide flexibility for learners and leverage the clustered nature of our target demographic.

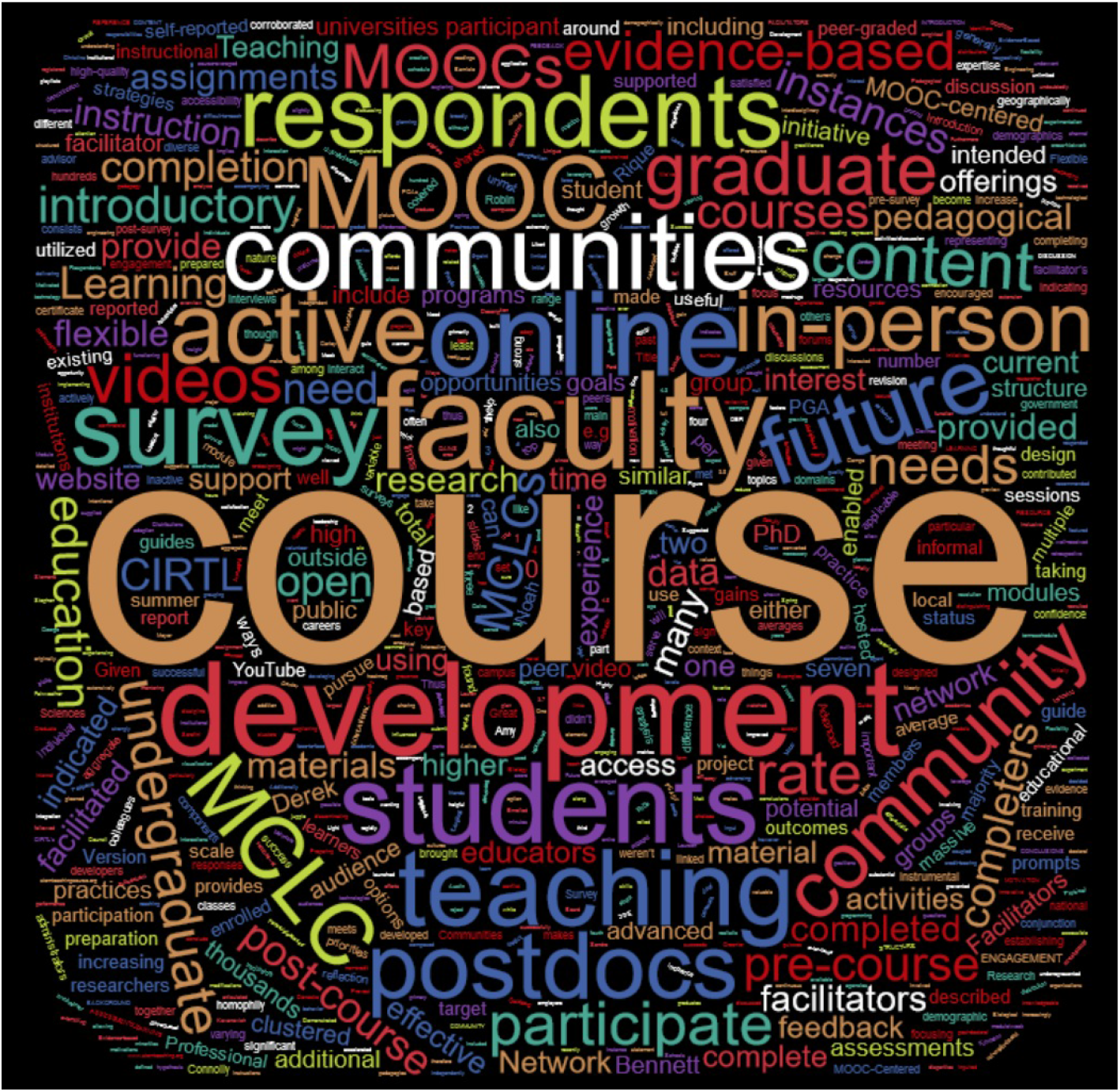

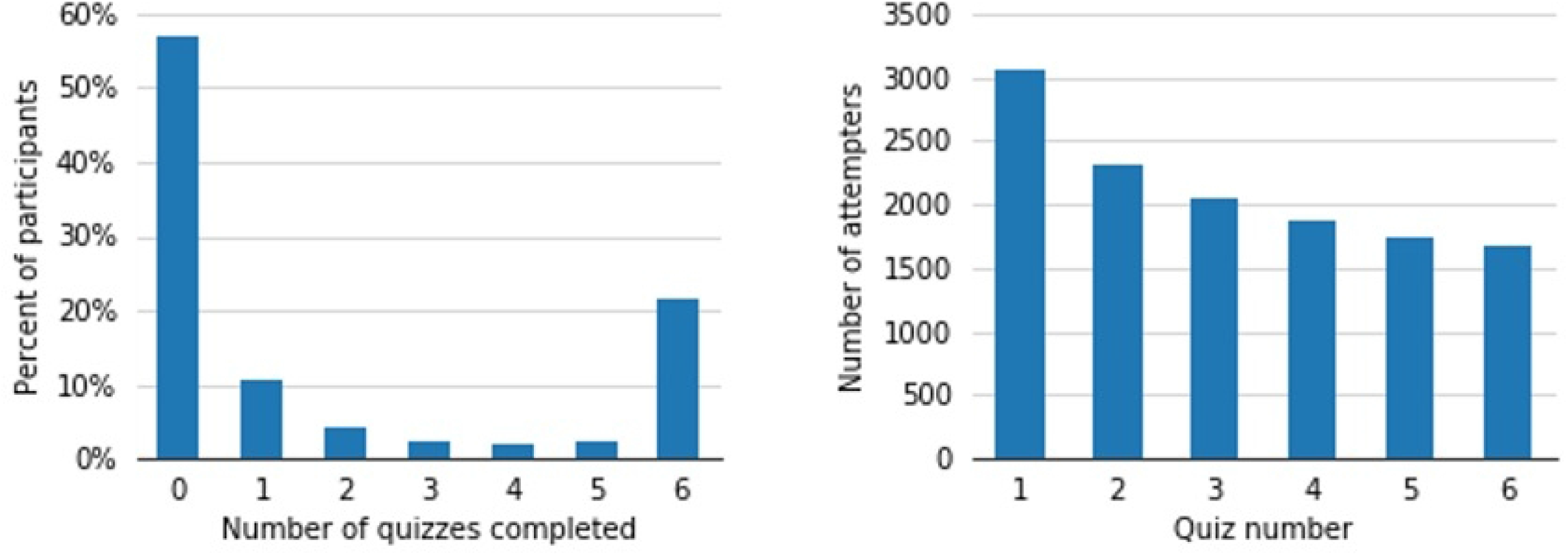

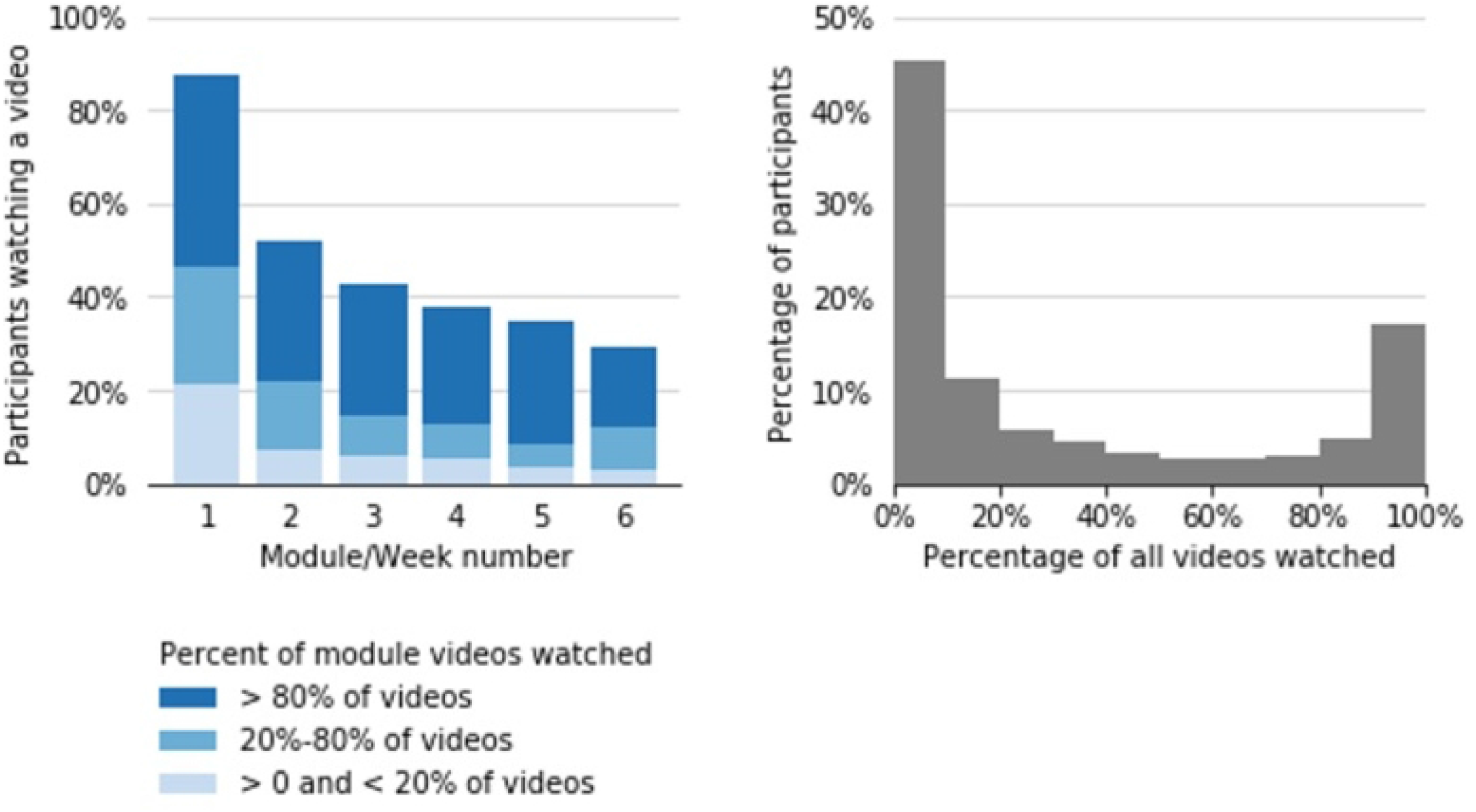

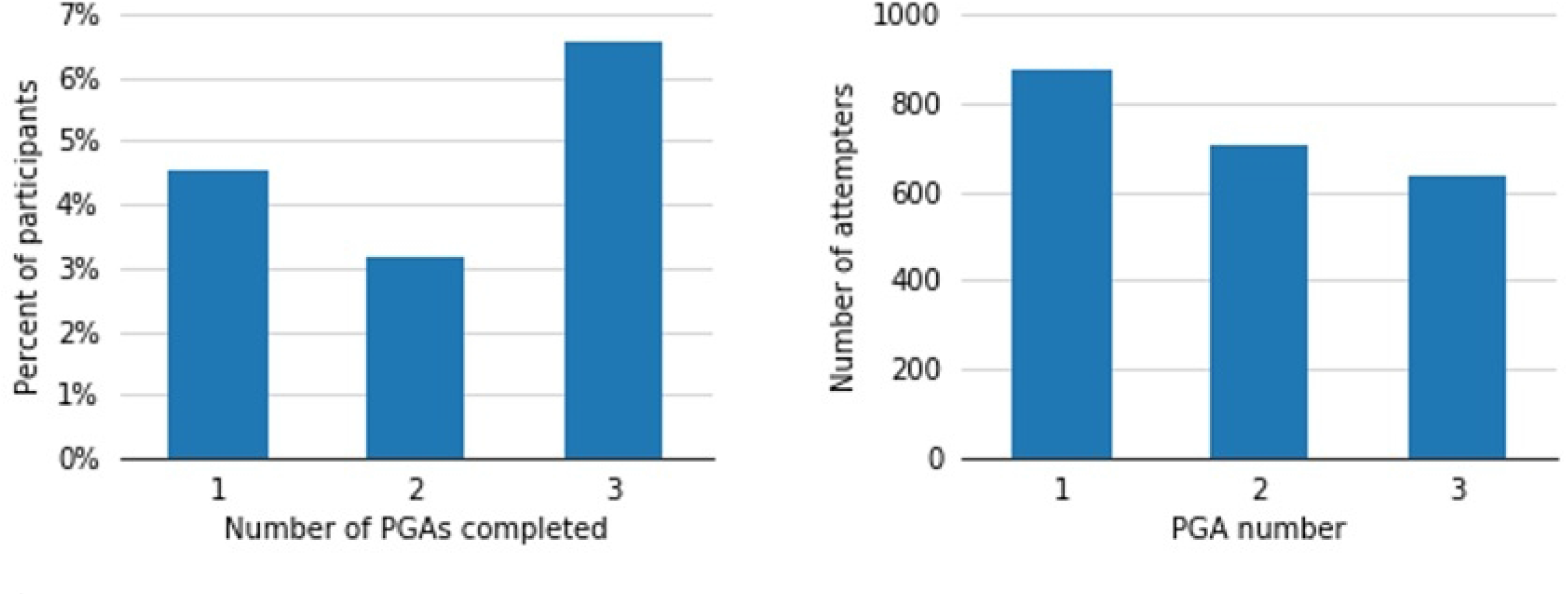

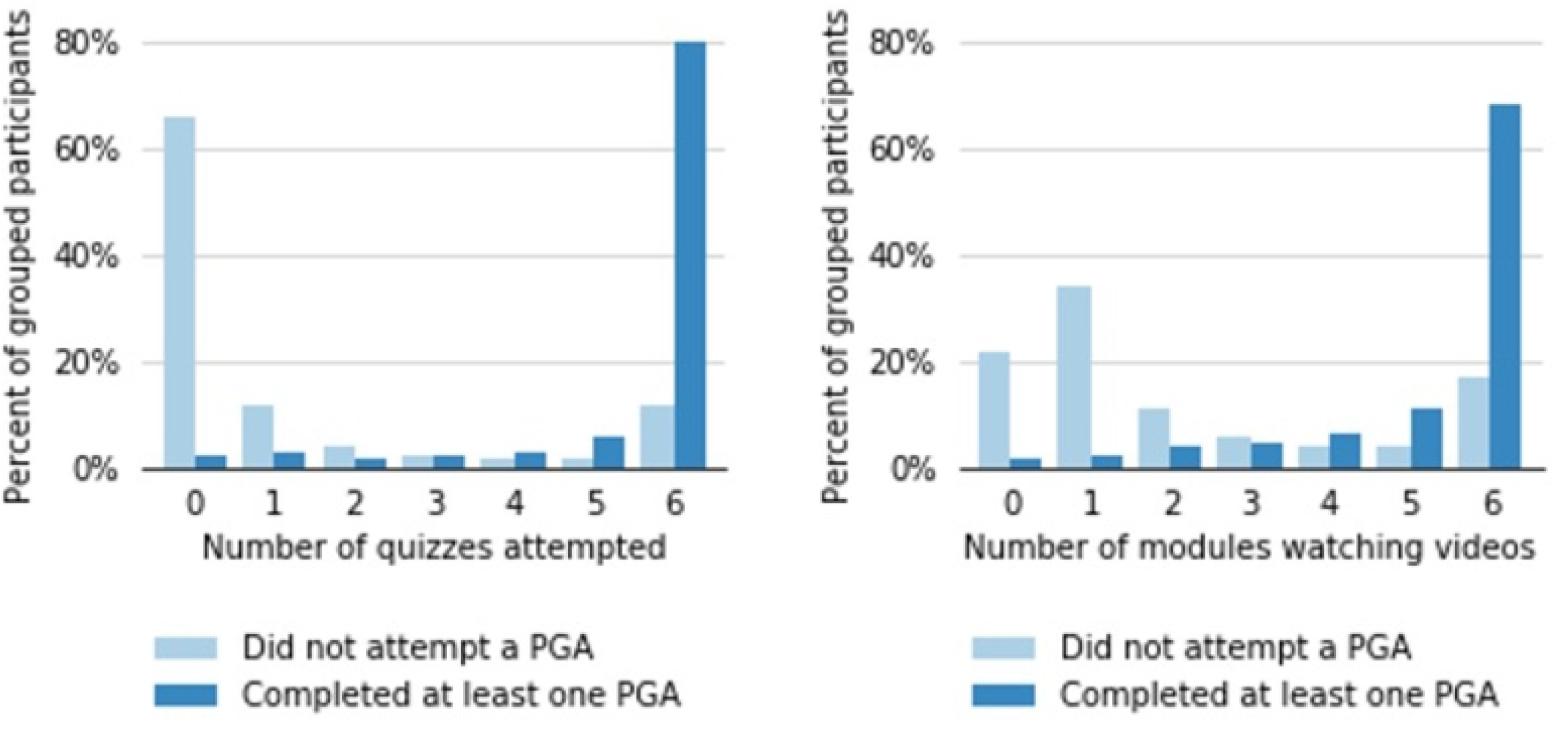

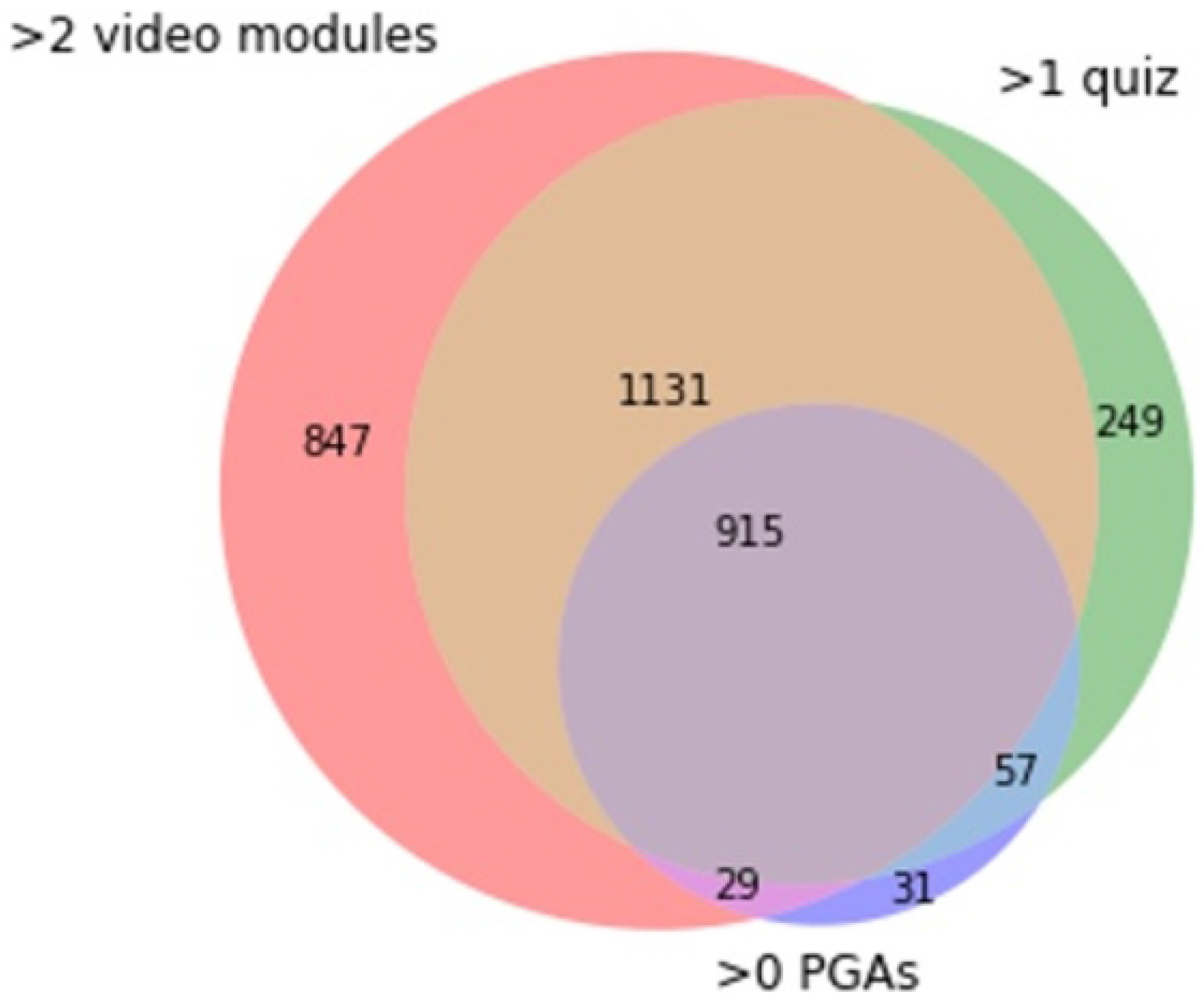

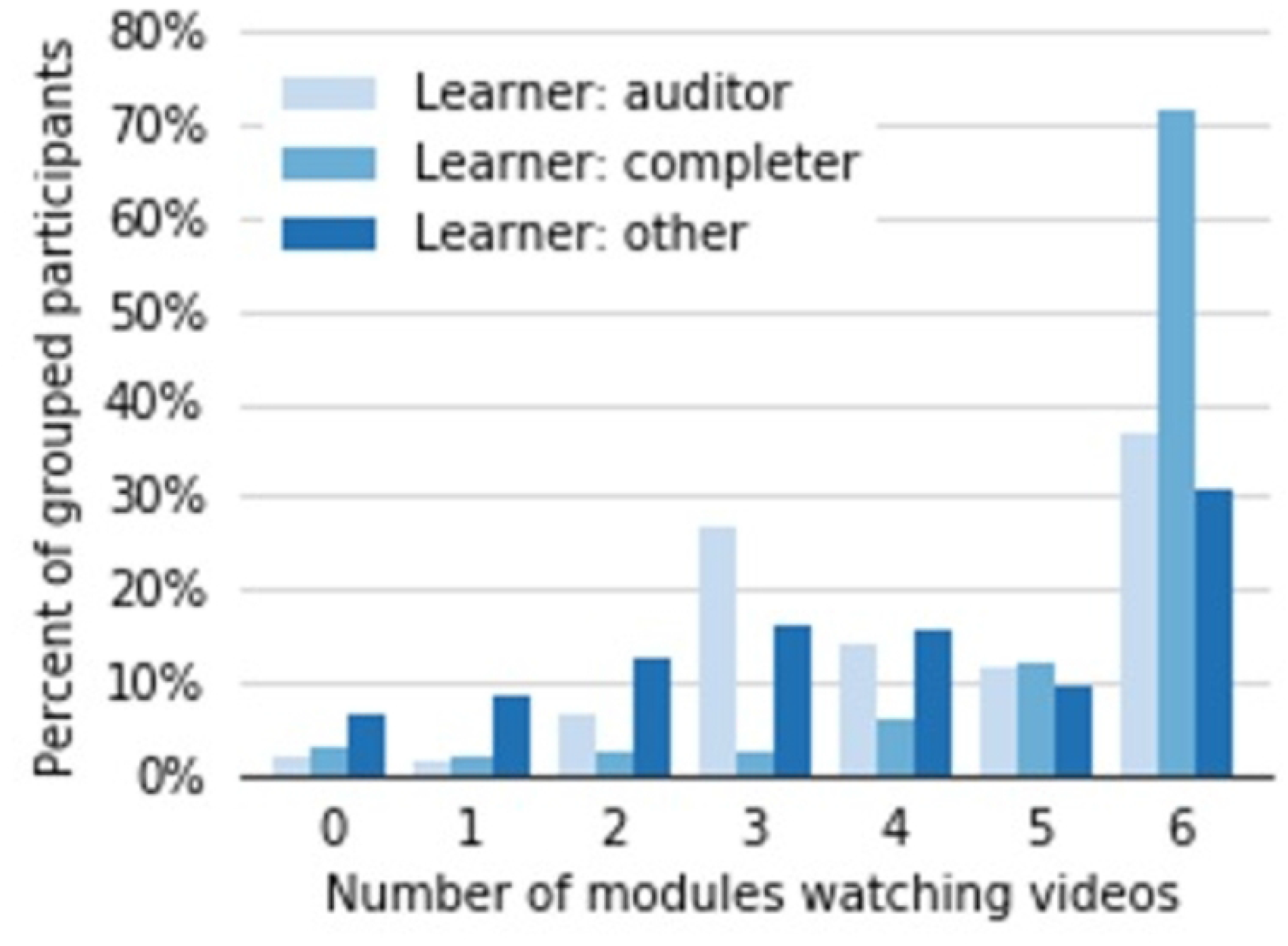

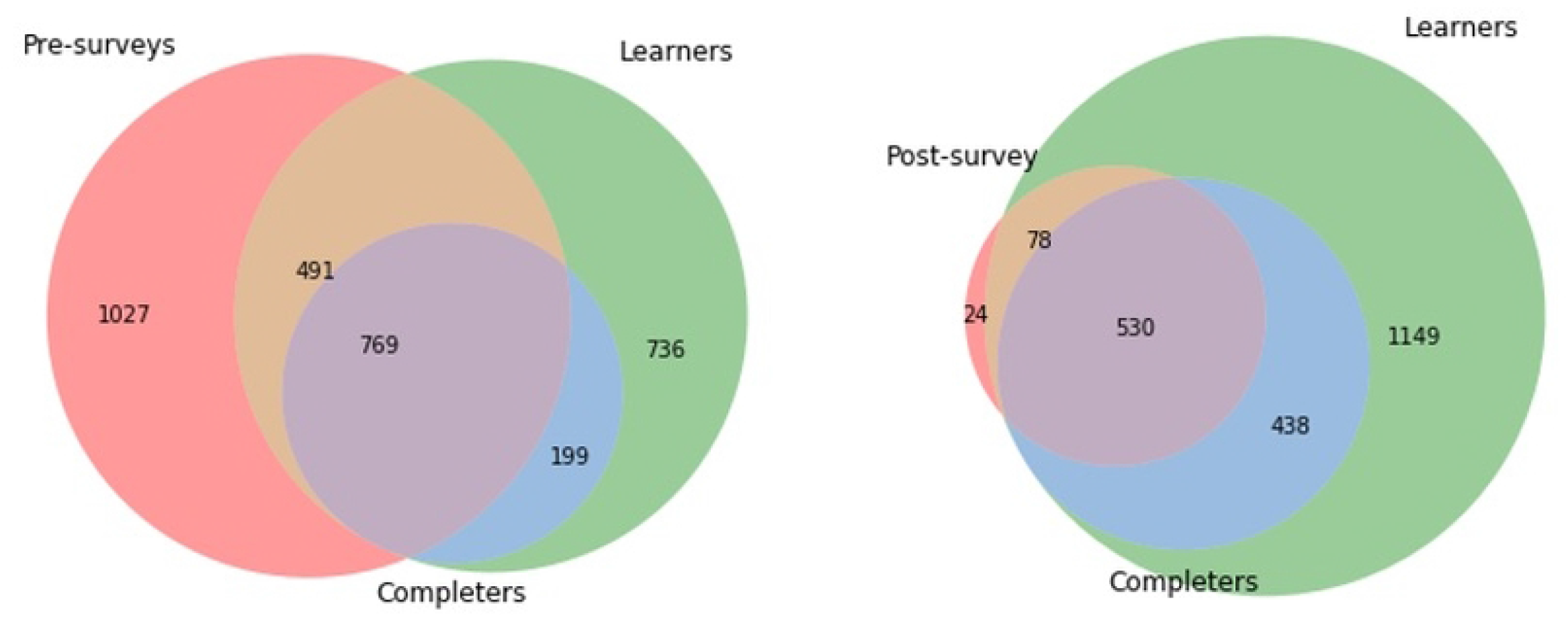

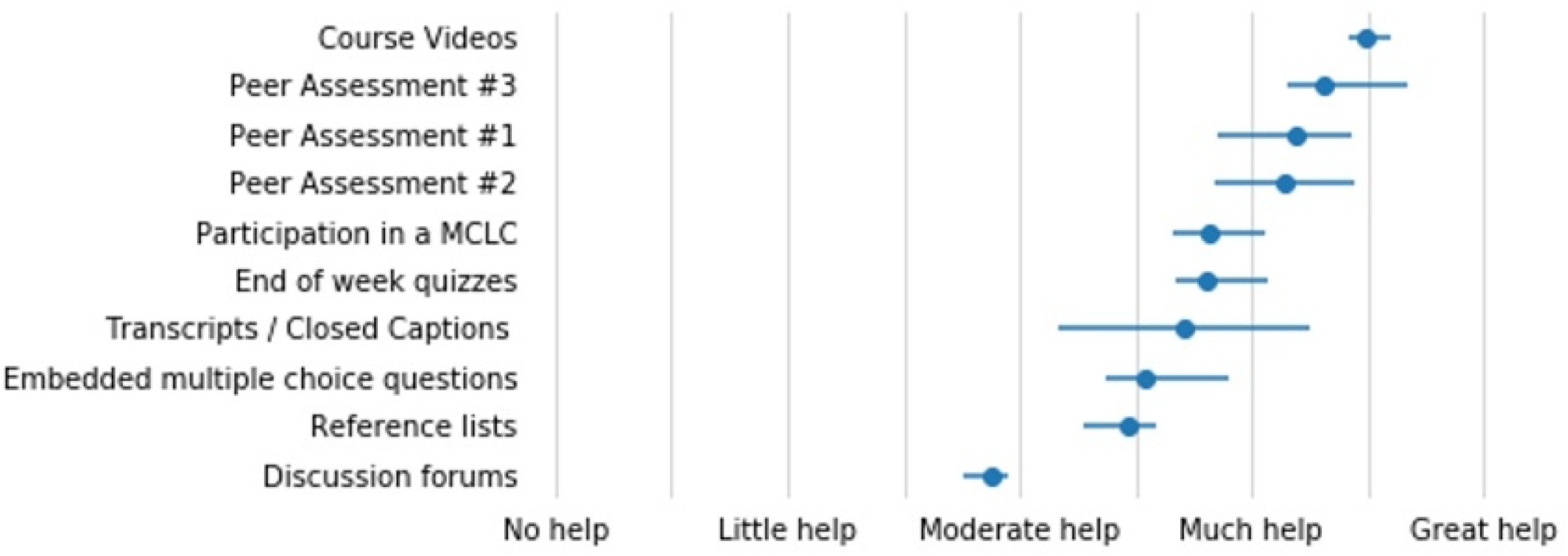

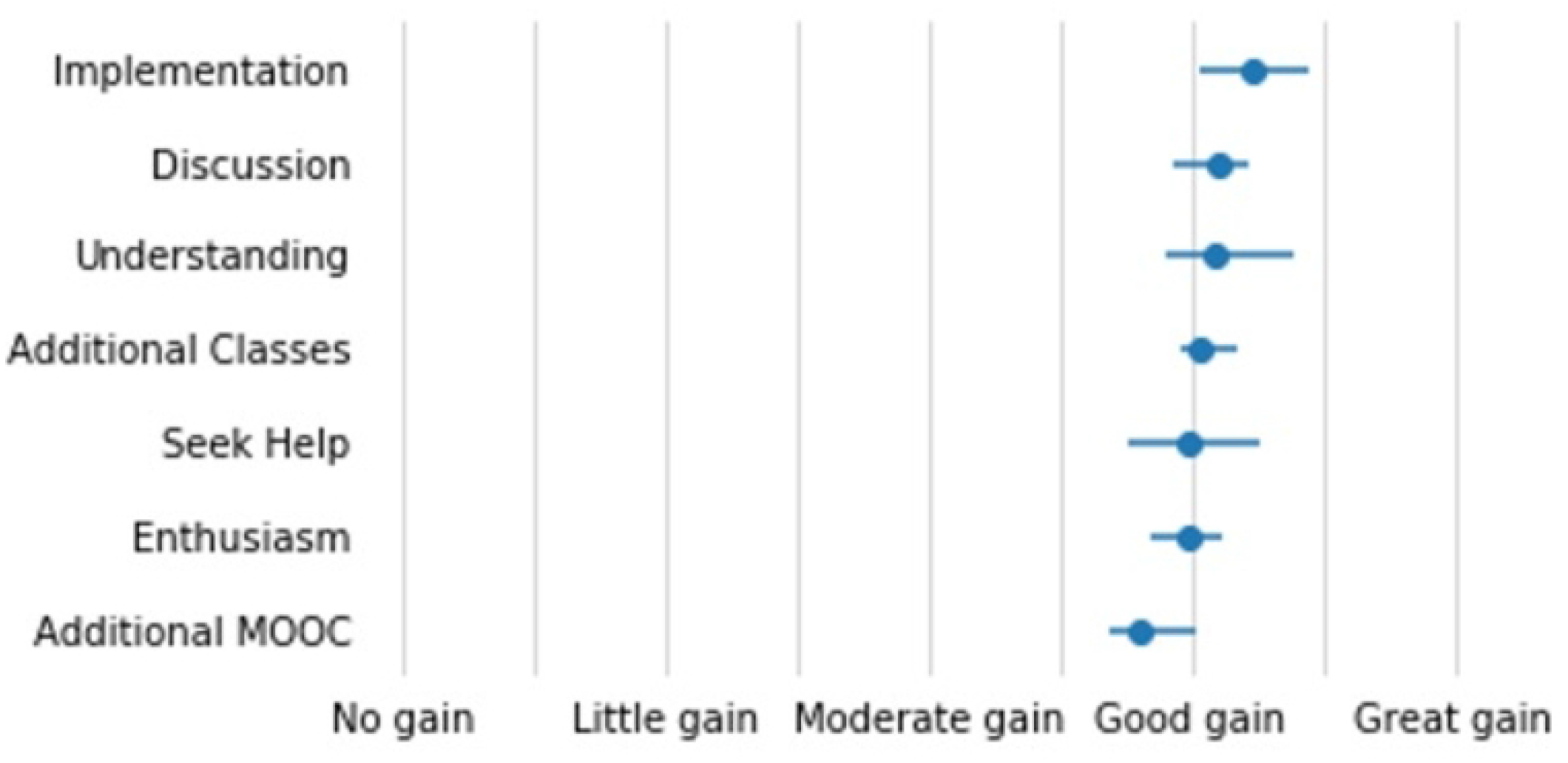

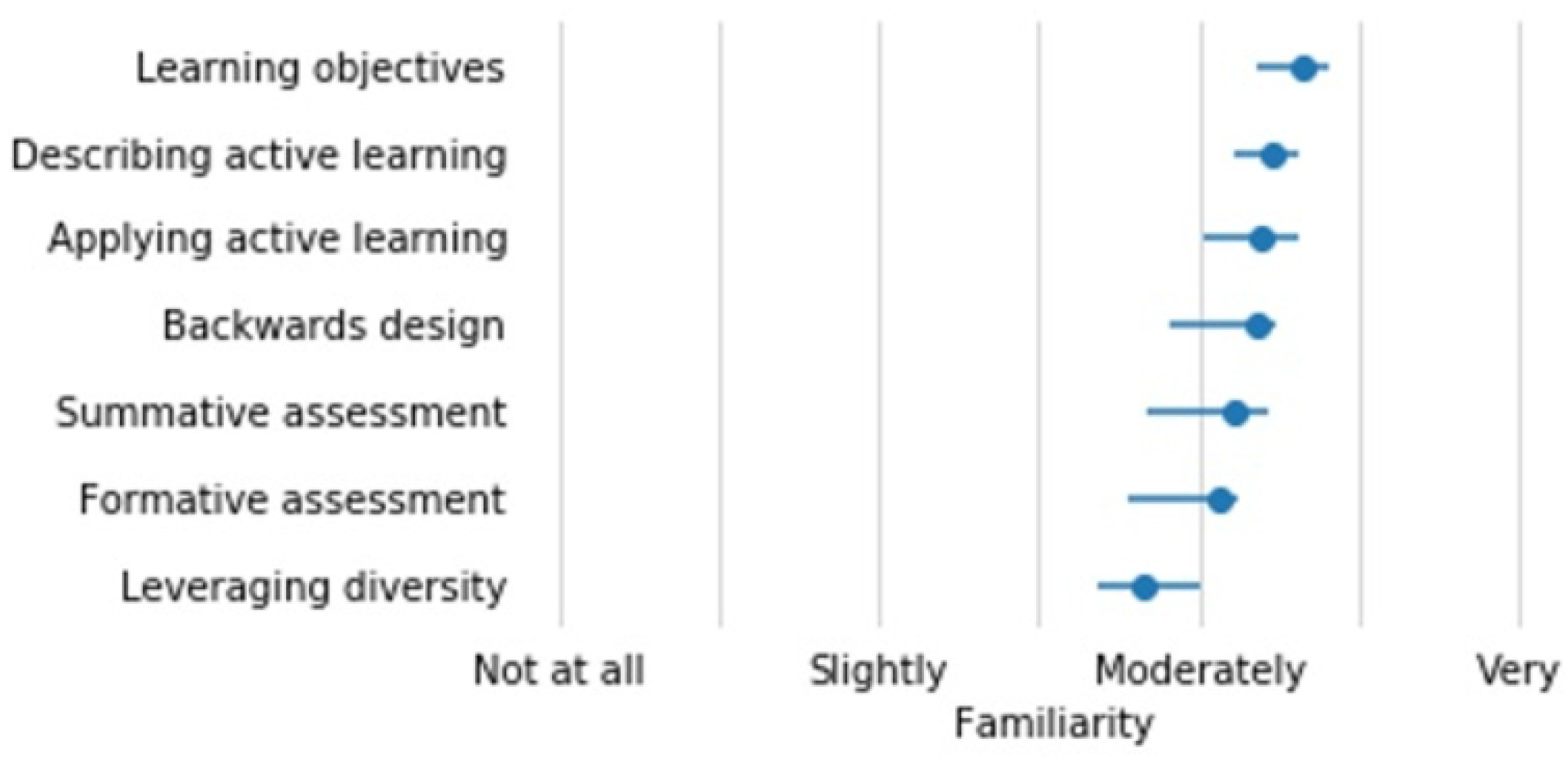

